# STL1267 alters HIF-1α and REV-ERBα localization and transcriptional responses in CoCl_2_-treated HeLa cells

**DOI:** 10.64898/2026.11.16.767081

**Authors:** Sangeet Makhija, Joshua D. Griffett, Kristine Griffett

## Abstract

**Background:** REV-ERB nuclear receptors regulate gene expression and cellular metabolism, but their role in the response to hypoxia-associated stress remains unclear. We examined how the REV-ERB agonist STL1267 affects protein distribution and gene expression in HeLa cells exposed to cobalt chloride (CoCl_2_).

**Methods and Results:** Immunofluorescence microscopy and RNA sequencing were used to compare vehicle-treated controls, cells exposed to CoCl_2_ plus vehicle, and cells exposed to CoCl_2_ plus STL1267. Representative images showed stronger nuclear HIF-1α staining after CoCl_2_ exposure. Nuclear staining was less pronounced with STL1267, while REV-ERBα showed more extranuclear staining. CoCl_2_ exposure altered 267 genes, and STL1267 treatment during CoCl_2_ exposure altered 132 genes at a false discovery rate below 0.05. STL1267 increased ATF4 expression alongside genes involved in amino-acid synthesis and one-carbon metabolism, including ASNS, PHGDH, PSAT1, PSPH and MTHFD2. CTH and CHAC1 also increased, highlighting changes in sulfur-amino-acid metabolism and glutathione turnover. Among the genes induced by CoCl_2_, CA9, BNIP3 and LDHA decreased with STL1267, whereas HK2, PDK1 and VEGFA did not change significantly.

**Conclusions:** STL1267 selectively reshapes hypoxia-associated gene expression and induces an ATF4-associated metabolic transcriptional program in CoCl_2_-treated HeLa cells, alongside qualitative changes in HIF-1α and REV-ERBα distribution. These findings identify a potential connection between protein compartmentalization and metabolic stress responses, providing a basis for investigating how REV-ERB ligands influence cellular adaptation to oxygen limitation.

## Introduction

Cells respond to low oxygen by changing the expression of genes involved in metabolism, survival and oxygen delivery. Much of this response is coordinated by hypoxia-inducible factor 1 (HIF-1), which consists of the oxygen-regulated HIF-1α subunit and its binding partner ARNT [1]. Oxygen-dependent prolyl hydroxylation promotes recognition of HIF-1α by the von Hippel–Lindau ubiquitin ligase complex, linking oxygen availability to HIF-1α degradation [2,3]. When oxygen becomes limiting, HIF-1α accumulates and supports the expression of genes that help cells respond to their changing environment.

Cobalt chloride (CoCl_2_) is widely used to induce hypoxia-like signaling in cultured cells because it stabilizes HIF proteins without requiring a reduction in the oxygen concentration of the culture environment [4]. This makes it useful for studying HIF-associated responses under defined treatment conditions. Its effects are not identical to those of low oxygen, however. Direct transcriptomic comparisons have shown differences between CoCl_2_ exposure and oxygen deprivation, so findings from this model need to be interpreted in the context of chemical pseudohypoxia [5].

REV-ERBα and REV-ERBβ are heme-binding nuclear receptors that repress gene expression and participate in circadian and metabolic regulation [6–8]. Their influence on metabolism extends beyond the core clock. In skeletal muscle, REV-ERBα regulates mitochondrial biogenesis, autophagy and oxidative capacity [9]. STL1267 is a synthetic REV-ERB agonist that binds the receptor and promotes recruitment of the nuclear receptor corepressor NCoR [10]. These properties provide a basis for examining how STL1267 affects cellular stress responses, including whether treatment changes the distribution of REV-ERBα and HIF-1α during CoCl_2_ exposure.

Another major regulator of stress adaptation is activating transcription factor 4 (ATF4). ATF4 coordinates genes involved in amino-acid availability and redox regulation and contributes to the response to mitochondrial stress [11,12]. In endothelial cells, Alhusban et al. showed that glucosamine induced an ATF4-dependent metabolic program that supported survival and angiogenesis under hypoxia and serum starvation [13]. Their findings raise a broader question: can other pharmacological treatments produce related changes in amino-acid metabolic gene expression during hypoxia-associated stress?

We investigated this question in CoCl_2_-treated HeLa cells. This model allows protein distribution and gene expression to be examined under the same treatment conditions in a readily manipulated cellular system. Tagged constructs made it possible to visualize HIF-1α and REV-ERBα, while RNA sequencing captured the broader transcriptional response. Comparing control cells with CoCl_2_-treated cells established the response to the chemical challenge; comparing CoCl_2_-treated cells with and without STL1267 then identified the additional effects of treatment. We focused on whether changes in protein staining patterns accompanied changes in hypoxia-associated transcripts and genes involved in amino-acid and one-carbon metabolism.

## Materials and methods

### Cell culture and treatments

HeLa cells were obtained from ATCC and maintained in EMEM (HyClone) supplemented with 10% fetal bovine serum (Corning) and 2.5 mM L-glutamine (Thermo Fisher Scientific). Cells were cultured at 37°C in a humidified incubator containing 5% CO_2_. Cell identity, mycoplasma status and passage information were documented upon receipt. Cells were assigned to three conditions: DMSO vehicle without CoCl_2_, CoCl_2_ plus DMSO, or CoCl_2_ plus STL1267. CoCl_2_ was used at 100 μM for 24 hours. In the STL1267 group, 10 μM STL1267 was added concurrently with CoCl_2_. The same treatment conditions were used for microscopy and RNA sequencing. Experiments were performed independently three times.

### Immunofluorescence microscopy

For imaging, HeLa cells were co-transfected with constructs encoding HA-tagged HIF-1α and Myc-tagged REV-ERBα before undergoing the treatments described above. Cells were fixed with 4% paraformaldehyde in PBS and blocked with normal goat serum. Tagged proteins were detected using anti-HA and anti-Myc primary antibodies from Thermo Fisher Scientific at 1:1,000, followed by secondary antibodies at 1:10,000. Nuclei were stained with DAPI. MitoID was used to label mitochondria in the HIF-1α images shown in Figure 1; the REV-ERBα images in Figure 2 contained REV-ERBα and DAPI staining. Images were acquired using an ECHO Revolve microscope and examined using ImageJ [14]. The microscopy was assessed qualitatively for differences in nuclear and extranuclear staining.

**Figure 1.**
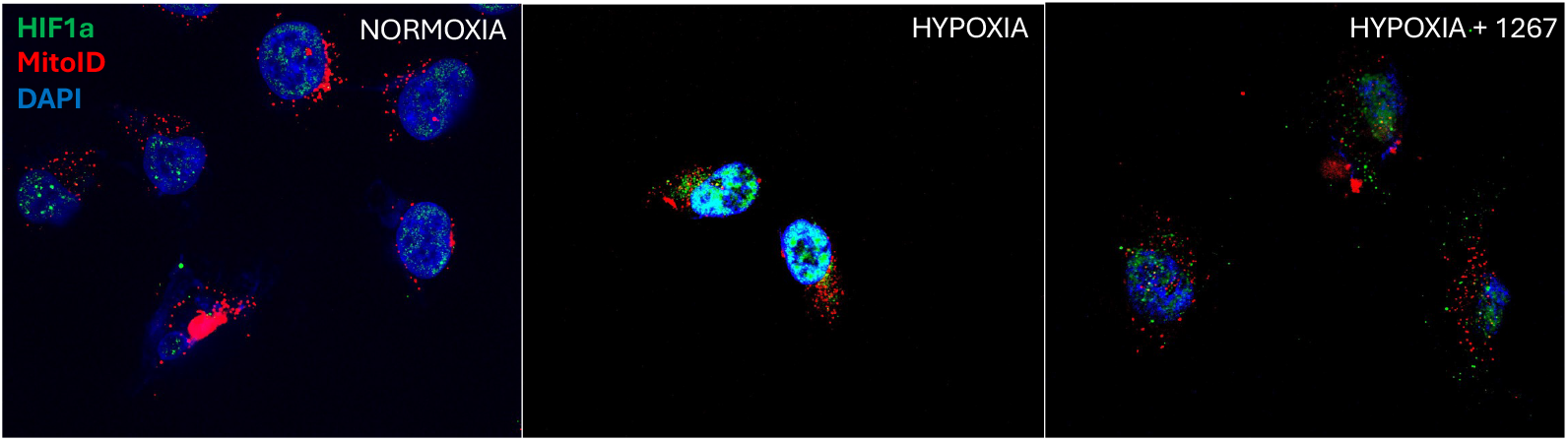
HIF-1α staining in CoCl_2_-treated HeLa cells with and without STL1267. Representative fluorescence images of HeLa cells treated for 24 hours with DMSO alone (left), 100 μM CoCl_2_ plus DMSO (middle), or 100 μM CoCl_2_ plus 10 μM STL1267 (right). HA-tagged HIF-1α was detected using an anti-HA antibody (green), mitochondria were labeled with MitoID (red), and nuclei were stained with DAPI (blue). Nuclear HIF-1α staining was more pronounced after CoCl_2_ exposure and appeared weaker in cells receiving STL1267. “Hypoxia” in the image labels refers to CoCl_2_-induced pseudohypoxia.

**Figure 2.**
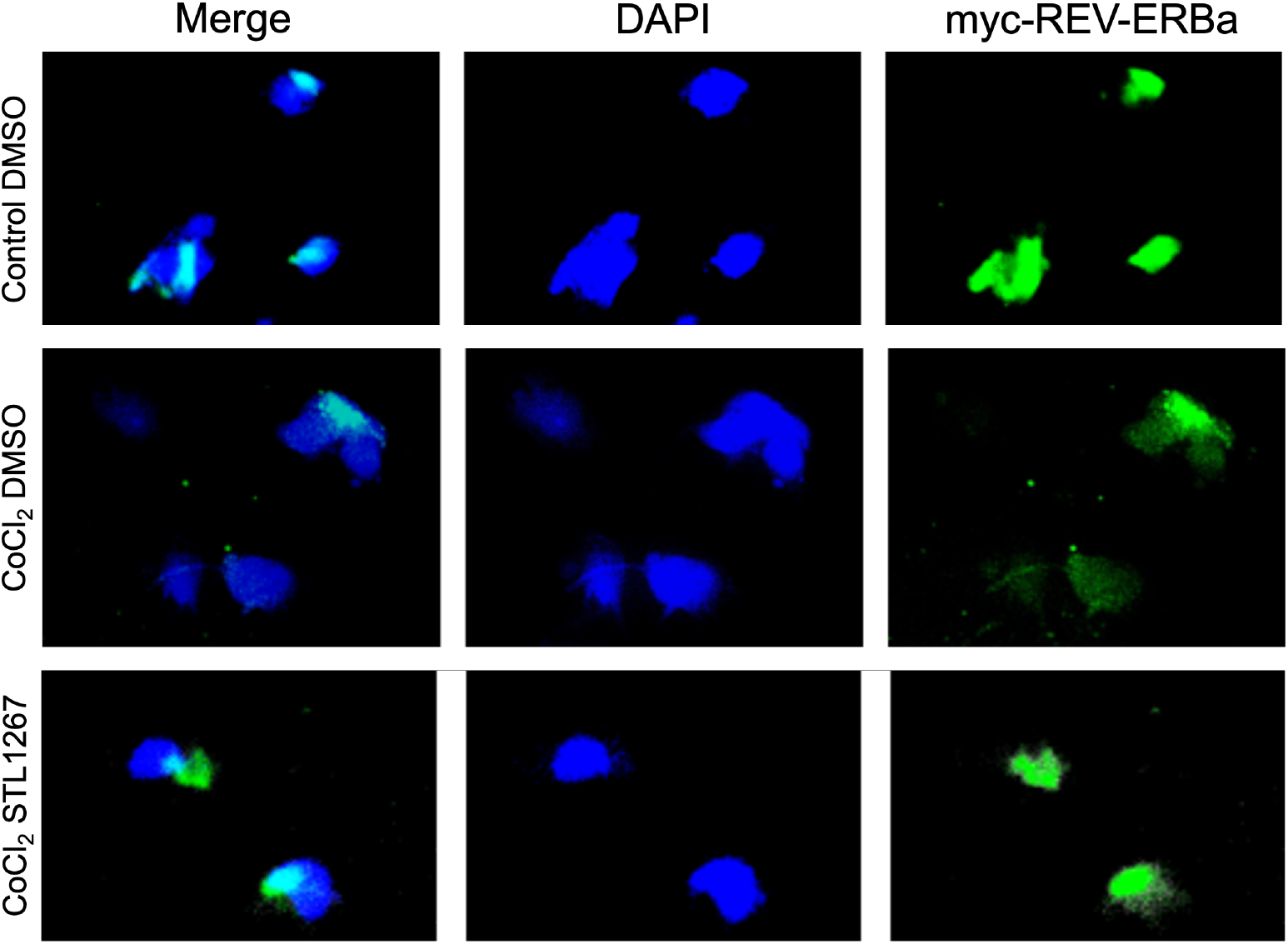
REV-ERBα staining shows an increased extranuclear distribution following STL1267 treatment. Representative fluorescence images of HeLa cells expressing Myc-tagged REV-ERBα after 24 hours of treatment with DMSO alone (top), 100 μM CoCl_2_ plus DMSO (middle), or 100 μM CoCl_2_ plus 10 μM STL1267 (bottom). REV-ERBα was detected using an anti-Myc antibody (green), and nuclei were stained with DAPI (blue). Columns show merged images (left), DAPI (middle), and REV-ERBα (right). Cyan in the merged images reflects overlap between the green and blue signals. Cells receiving STL1267 showed more REV-ERBα staining outside the DAPI-defined nuclear region.

### RNA sequencing and differential-expression analysis

After treatment, cells were collected, counted and placed in Zymo RNA preservation reagent according to Plasmidsaurus sample submission guidelines. Library preparation, sequencing and data analysis were performed by Plasmidsaurus. Two comparisons were evaluated: CoCl_2_ plus vehicle relative to control without CoCl_2_, and CoCl_2_ plus STL1267 relative to CoCl_2_ plus vehicle. Positive reported logFC values indicate higher expression in the first condition named in each comparison. Genes with a false discovery rate (FDR) below 0.05 were considered differentially expressed. The two comparisons were interpreted separately. Selected genes were examined to describe changes in hypoxia-associated transcription, amino-acid metabolism, serine synthesis and one-carbon metabolism. This gene selection was exploratory; a formal pathway-enrichment analysis was not performed. Reported logFC values were retained as supplied in the differential-expression results.

## Results and discussion

CoCl_2_ exposure produced a clear change in HIF-1α staining. Compared with DMSO-treated controls, CoCl_2_-treated cells showed stronger HA-tagged HIF-1α signal within DAPI-labeled nuclei. Nuclear staining was less pronounced when STL1267 was added concurrently with CoCl_2_, with some signal remaining outside the nucleus (**Fig. 1**). This difference suggested that STL1267 affected nuclear HIF-1α accumulation during the chemical challenge and led us to examine whether hypoxia-associated gene expression also changed.

REV-ERBα showed a corresponding difference in its staining pattern. Myc-tagged REV-ERBα was predominantly nuclear in control and CoCl_2_-treated cells, whereas cells receiving CoCl_2_ and STL1267 showed more signal outside the DAPI-defined nuclear region (**Fig. 2**). The images therefore suggest changes in the distribution of both tagged proteins under the same treatment conditions: less nuclear HIF-1α staining and more extranuclear REV-ERBα staining. Because Figure 2 contains no organelle marker, the extranuclear REV-ERBα signal cannot be assigned to a specific compartment. Such an assignment would require additional labeling and quantitative spatial analysis [15].

These observations raise a possible connection between protein distribution and transcription. Reduced nuclear HIF-1α could limit its contribution to hypoxia-responsive gene expression, while a shift in REV-ERBα distribution could change its access to nuclear targets. The two responses might be linked, or they could reflect separate effects of STL1267. We therefore examined the RNA-seq results to determine which transcriptional changes accompanied the imaging observations. CoCl_2_ exposure altered 267 of the 14,861 genes included in the comparison at FDR < 0.05, with 213 increased and 54 decreased genes (**Fig. 3A**). Several genes associated with the hypoxic response increased, including CA9, BNIP3, PDK1, HK2, VEGFA and LDHA (**Table 1; Fig. 3B**). CA9 showed the largest increase among these selected genes, with a reported logFC of +5.56, followed by BNIP3 (+2.38) and PDK1 (+2.03). PFKFB3 and G6PD also increased. This expression pattern, together with the stronger nuclear HIF-1α staining, was consistent with induction of a hypoxia-associated response by CoCl_2_.

**Table 1:** Selected gene expression changes in the two RNA-seq comparisons.

| Gene | CoCl<sub>2</sub><br>logFC | CoCl<sub>2</sub><br>FDR | STL1267<br>logFC | STL1267<br>FDR |
|---|---|---|---|---|
| CA9 | +5.561 | 5.98e-22 | -0.950 | 1.02e-09 |
| BNIP3 | +2.381 | 6.72e-21 | -0.437 | 3.87e-05 |
| HK2 | +1.298 | 2.02e-09 | -0.015 | 0.978 |
| LDHA | +0.833 | 4.19e-12 | -0.225 | 0.0378 |
| PDK1 | +2.034 | 2.65e-14 | -0.113 | 0.7 |
| VEGFA | +0.952 | 1.56e-08 | +0.227 | 0.256 |
| PFKFB3 | +0.711 | 0.00108 | -0.156 | 0.73 |
| G6PD | +0.322 | 0.000237 | -0.010 | 0.973 |
| ATF4 | +0.114 | 0.393 | +0.921 | 9.25e-12 |
| ASNS | -0.369 | 0.102 | +1.950 | 4.57e-14 |
| PHGDH | -0.094 | 0.564 | +0.878 | 1.52e-10 |
| PSAT1 | -0.309 | 0.00161 | +1.263 | 4.57e-14 |
| PSPH | -0.092 | 0.755 | +0.787 | 1.24e-06 |
| MTHFD2 | -0.142 | 0.37 | +0.702 | 4.53e-08 |
| CTH | -0.157 | 0.839 | +1.616 | 1.1e-06 |
| CHAC1 | -0.075 | 0.969 | +2.652 | 4.66e-08 |
| NUPR1 | -0.294 | 0.587 | +2.166 | 1.52e-10 |
| CBS | -0.270 | 0.172 | +0.594 | 0.000236 |
| SLC7A11 | +0.171 | 0.139 | +0.543 | 3.52e-07 |
| OGT | -0.042 | 0.878 | +0.356 | 0.00424 |

CoCl<sub>2</sub>: CoCl<sub>2</sub> plus vehicle relative to control without CoCl<sub>2</sub>. STL1267: CoCl<sub>2</sub> plus STL1267 relative to CoCl<sub>2</sub> plus vehicle. Values are reported logFC and FDR; FDR < 0.05 defines differential expression.

**Figure 3.**
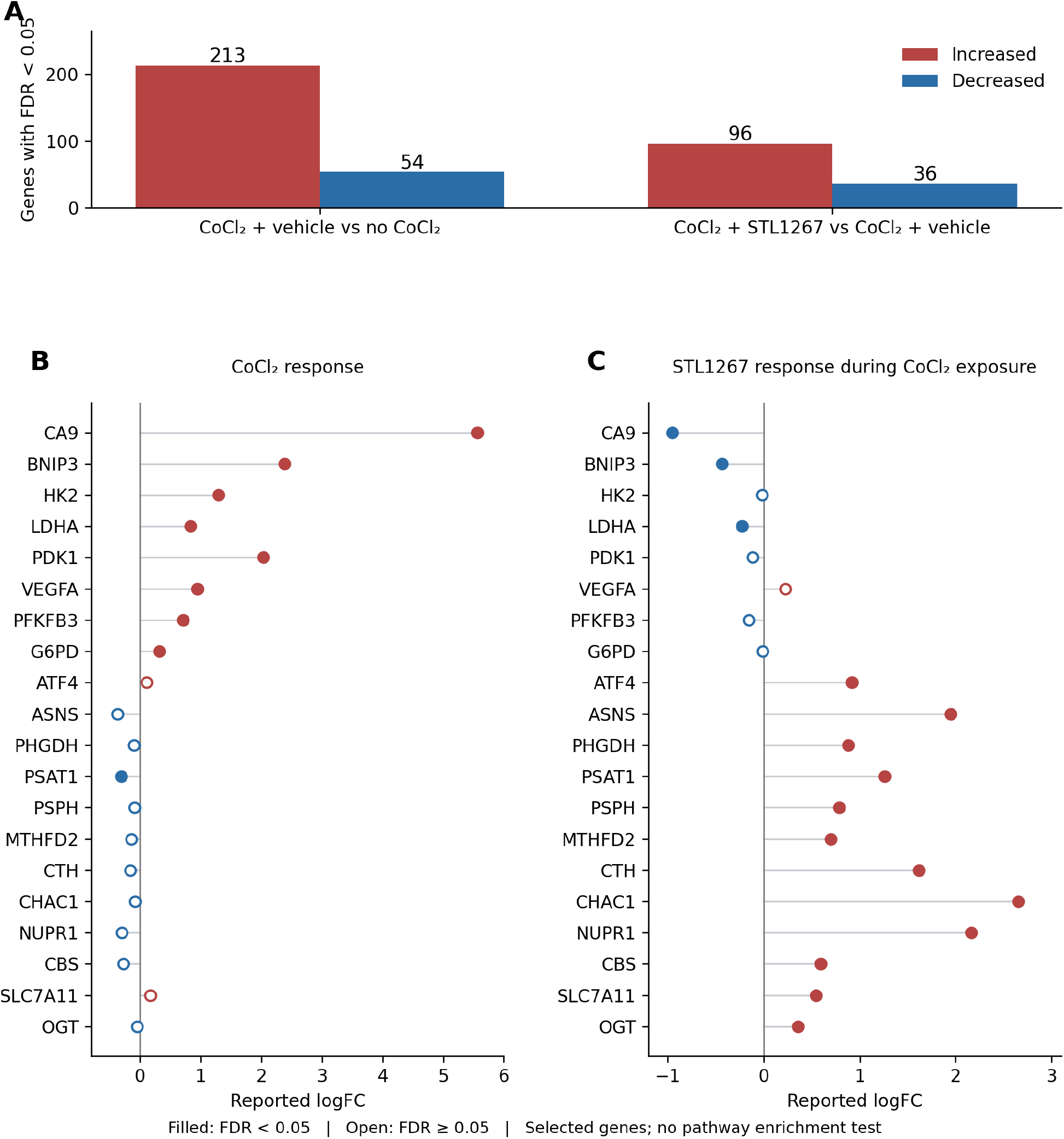
Transcriptional responses to CoCl_2_ and STL1267 in HeLa cells. (A) Numbers of differentially expressed genes at FDR < 0.05. CoCl_2_ plus vehicle relative to control without CoCl_2_ yielded 267 differentially expressed genes, comprising 213 increased and 54 decreased genes. CoCl_2_ plus STL1267 relative to CoCl_2_ plus vehicle yielded 132 differentially expressed genes, comprising 96 increased and 36 decreased genes. (B, C) Reported logFC values for the selected genes listed in Table 1 in the CoCl_2_ and STL1267 comparisons, respectively. Red indicates positive logFC and blue indicates negative logFC. Filled points indicate FDR < 0.05; open points indicate FDR ≥ 0.05. Plots show differential-expression estimates for selected genes.

Adding STL1267 during CoCl_2_ exposure altered 132 of the 14,883 genes included in the treatment comparison: 96 increased and 36 decreased (**Fig. 3A**). CA9, BNIP3 and LDHA, which had increased with CoCl_2_, decreased with STL1267. Their reported treatment-associated logFC values were −0.95, −0.44 and −0.23, respectively, with FDR < 0.05 for each gene. These changes accompanied the weaker nuclear HIF-1α staining in Figure 1. HK2, PDK1, VEGFA, PFKFB3 and G6PD did not change significantly with STL1267, indicating that treatment affected selected components of the CoCl_2_ response. The opposite directions observed for CA9, BNIP3 and LDHA in the two comparisons do not, by themselves, establish a return to control expression levels.

The treatment response also extended beyond hypoxia-associated genes. ATF4 increased with STL1267 (logFC +0.92; FDR 9.25 × 10^−12^), along with ASNS (+1.95), PHGDH (+0.88), PSAT1 (+1.26), PSPH (+0.79) and MTHFD2 (+0.70) (**Table 1; Fig. 3C**). The coordinated increase in transcripts encoding PHGDH, PSAT1 and PSPH, the three enzymes of de novo serine synthesis, was a prominent feature. This pattern differed from the response to CoCl_2_ alone, in which ATF4, PHGDH, PSPH and MTHFD2 did not change significantly and PSAT1 decreased (logFC −0.31; FDR 0.0016). STL1267 thus produced an additional metabolic expression pattern during CoCl_2_ exposure.

The increase in ATF4 and related metabolic genes fits with its established role in cellular stress responses. ATF4 helps coordinate amino-acid availability and resistance to oxidative stress, and Quirós et al. identified it as a major regulator of the response to mitochondrial perturbation [11,12]. ATF4 is also regulated at the level of translation. In HeLa cells exposed to low oxygen, Blais et al. found increased ATF4 translation without a corresponding increase in total ATF4 mRNA [16]. This distinction matters here: the RNA-seq results identify an ATF4-associated expression pattern, while measurements of ATF4 protein and its dependence on upstream stress signaling would help establish how that pattern develops.

The serine and one-carbon metabolic genes provide a more specific direction for investigating this response. Serine synthesis supports several biosynthetic demands in cancer cells, and studies in breast and lung cancer have linked its regulation to cellular growth and stress-associated transcription [17,18]. Serine can also supply one-carbon units for nucleotide synthesis, connecting amino-acid metabolism to the demands of proliferating cells [19]. The increase in MTHFD2 alongside PHGDH, PSAT1 and PSPH points to this broader metabolic network, which includes both cytosolic and mitochondrial folate pathways [20]. In the present study, these changes describe transcript abundance; whether they increase serine synthesis or one-carbon flux remains to be tested.

The overlap with the endothelial findings of Alhusban et al. is particularly relevant. Their study showed that glucosamine induced ATF4-dependent amino-acid metabolism and supported endothelial survival and angiogenesis under hypoxia and serum starvation [13]. ASNS, PHGDH, PSAT1, PSPH and MTHFD2 were among the expression components shared with our STL1267 response. This overlap suggests that different treatments can engage related metabolic transcriptional programs. It also provides a practical starting point for testing STL1267 in endothelial cells, where changes in gene expression can be evaluated alongside survival, barrier function and angiogenic behavior. Whether STL1267 produces similar functional effects will depend on experiments in those cells.

Sulfur-amino-acid metabolism and glutathione turnover were another feature of the treatment response. CTH (+1.62), CBS (+0.59) and SLC7A11 (+0.54) increased with STL1267. CBS and CTH participate in trans-sulfuration, and stress-responsive induction of CTH has previously been linked to changes in cellular thiol metabolism [21]. SLC7A11 encodes the light-chain component of the cystine/glutamate antiporter, connecting extracellular cystine availability to intracellular cysteine supply [22]. Their increased expression suggests that cysteine handling is part of the broader metabolic response to treatment.

CHAC1 showed an especially strong increase (logFC +2.65). CHAC1 encodes a glutathione-degrading enzyme whose expression is regulated by ATF4 and other stress-responsive transcription factors [23]. Its induction alongside genes involved in cysteine supply makes the overall effect on glutathione difficult to predict from RNA alone. The expression pattern could accompany changes in both substrate availability and glutathione breakdown. Measurements of reduced and oxidized glutathione, oxidative stress and cell viability would establish whether this response helps maintain redox balance or accompanies cellular injury. NUPR1 also increased (+2.17), adding to the evidence for a broader stress-associated transcriptional response. The imaging and RNA-seq results identify a common set of questions about how STL1267 acts during CoCl_2_ exposure. Does the change in REV-ERBα distribution influence HIF-1α accumulation? Does reduced nuclear HIF-1α contribute to the selective decrease in hypoxia-associated transcripts? And is the ATF4-associated response connected to either localization change? A time course combining protein measurements with gene expression would help establish the order of these events. Measurements of total protein and nuclear and extranuclear fractions would distinguish redistribution from changes in abundance, while perturbation of NR1D1, NR1D2 and ATF4 would test their contributions to the response.

The scope of the current findings is defined by the experimental model and the measurements available. The microscopy is qualitative and examines transfected, tagged proteins, so quantitative imaging and confirmation of endogenous protein distribution are needed. The RNA-seq comparisons describe separate responses to CoCl_2_ and to STL1267 during CoCl_2_ exposure; an STL1267-only condition would be needed to test whether treatment effects depend on the chemical challenge. CoCl_2_ also differs from reduced oxygen [4,5]. Testing the response under controlled low oxygen and in additional cell types will therefore be necessary to determine how broadly these observations apply. Within this model, STL1267 was accompanied by changes in HIF-1α and REV-ERBα staining patterns and by a transcriptional response that combined selective decreases in hypoxia-associated genes with increased expression of amino-acid metabolic genes. These findings provide a focused basis for examining how protein distribution and metabolic gene regulation contribute to the cellular response to STL1267.

## Conclusion

STL1267 produced a distinct response in CoCl_2_-treated HeLa cells, combining lower expression of selected hypoxia-associated genes with increased expression of genes involved in amino-acid synthesis, serine biosynthesis and one-carbon metabolism. These transcriptional changes accompanied an apparent reduction in nuclear HIF-1α staining and a more extranuclear distribution of REV-ERBα. The contribution of this study lies in bringing these observations together: STL1267-associated changes in protein distribution occur alongside a metabolic expression program that differs from the response to CoCl_2_ alone. These findings raise a broader question about how cells coordinate hypoxia-associated signaling with metabolic stress responses. Protein compartmentalization may be one point of connection between these processes. Establishing whether REV-ERBα redistribution contributes to changes in HIF-1α and ATF4-associated transcription could extend our understanding of REV-ERB biology beyond its established role in nuclear transcriptional repression. It would also help clarify how pharmacological modulation of these receptors influences the balance among different stress-response programs.

This work provides a starting point for connecting circadian nuclear receptor biology, hypoxia signaling and metabolic adaptation. Quantitative localization studies, receptor perturbation and experiments under controlled low oxygen can establish which parts of the response are mechanistically linked. Extending these studies to endothelial and skeletal muscle cells would reveal whether the same transcriptional program supports vascular or muscle function during oxygen limitation, while studies in additional cancer models could determine whether it contributes to tumor-cell adaptation or vulnerability. Defining these cell-specific consequences is an important step toward understanding where REV-ERB-directed treatments may have therapeutic value.

## Supporting information

Image analysis data

RNAseq data

RNAseq data

GSEA

Enrichment

## Declarations

### Ethics approval

This study used the established human HeLa cell line obtained from ATCC. No human participants were recruited, and no human tissue samples were collected. Institutional ethics approval and informed consent to participate were not required for these cell-line experiments.

### Data availability

All sequencing data will be deposited into NCBI GEO per NIH guidelines.

### Author contributions

S.M. – Conceptualization and Investigation of the project; Formal Analysis; Validation; Writing – review and editing.

J.D.G. – Investigation; Validation; Writing – review and editing.

K.G. – Conceptualization and Investigation of the project; Formal Analysis; Validation; Resources; Funding Acquisition; Writing – original draft; Writing – review and editing.

## Acknowledgments

We thank Plasmidsaurus for providing RNAseq data and analysis.

## Use of artificial intelligence

Generative AI was used to assist with manuscript drafting and language editing. The authors are responsible for the accuracy of the scientific content, interpretation of the results and final wording.

