## Supplementary material for "STL1267 alters HIF-1α and REV-ERBα localization and transcriptional responses in CoCl_2_-treated HeLa cells": Image analysis data

Figure 1

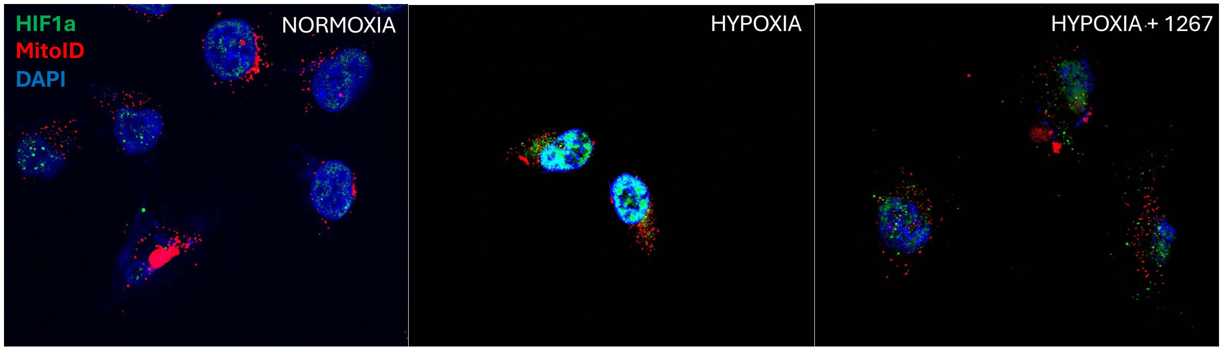

### Figure 1 – Masking and quantification

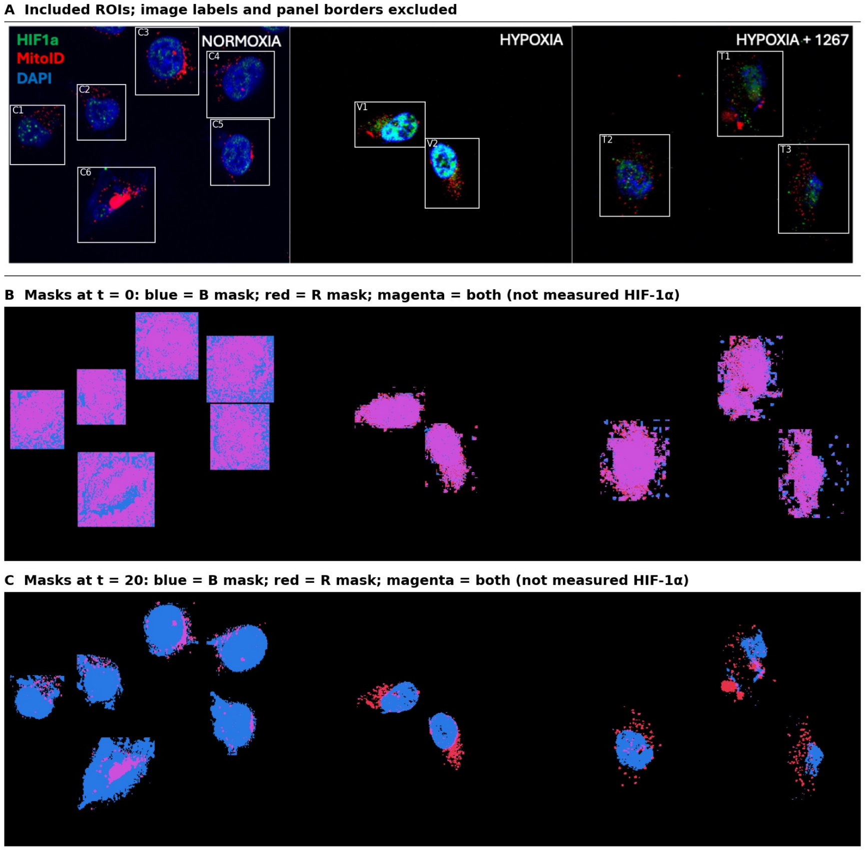


| Measurement | Control | $CoCl_2$ + vehicle | $CoCl_2$ + STL1267 |
|---|---|---|---|
| **Blue:** DAPI channel | 321,283 | 274,412 | 114,219 |
| **Green (HIF1a):** green outside blue mask | 0 | 763 | 926 |
| **Cyan:** green inside blue mask | 37,870 | 210,940 | 65,545 |
| **Red (Mitochondria):** MitoID channel | 60,892 | 63,685 | 65,103 |
| **Yellow:** green overlapping red mask | 36,826 | 208,963 | 65,182 |

| Panel | Inside blue mask | Outside blue mask | Overlapping red mask |
|---|---|---|---|
| Control | 94.2% | 5.8% | 10.5% |
| $CoCl_2$ + vehicle | 89.1% | 10.9% | 11.9% |
| $CoCl_2$ + STL1267 | 58.3% | 41.7% | 18.5% |

Figure 1-analyzed

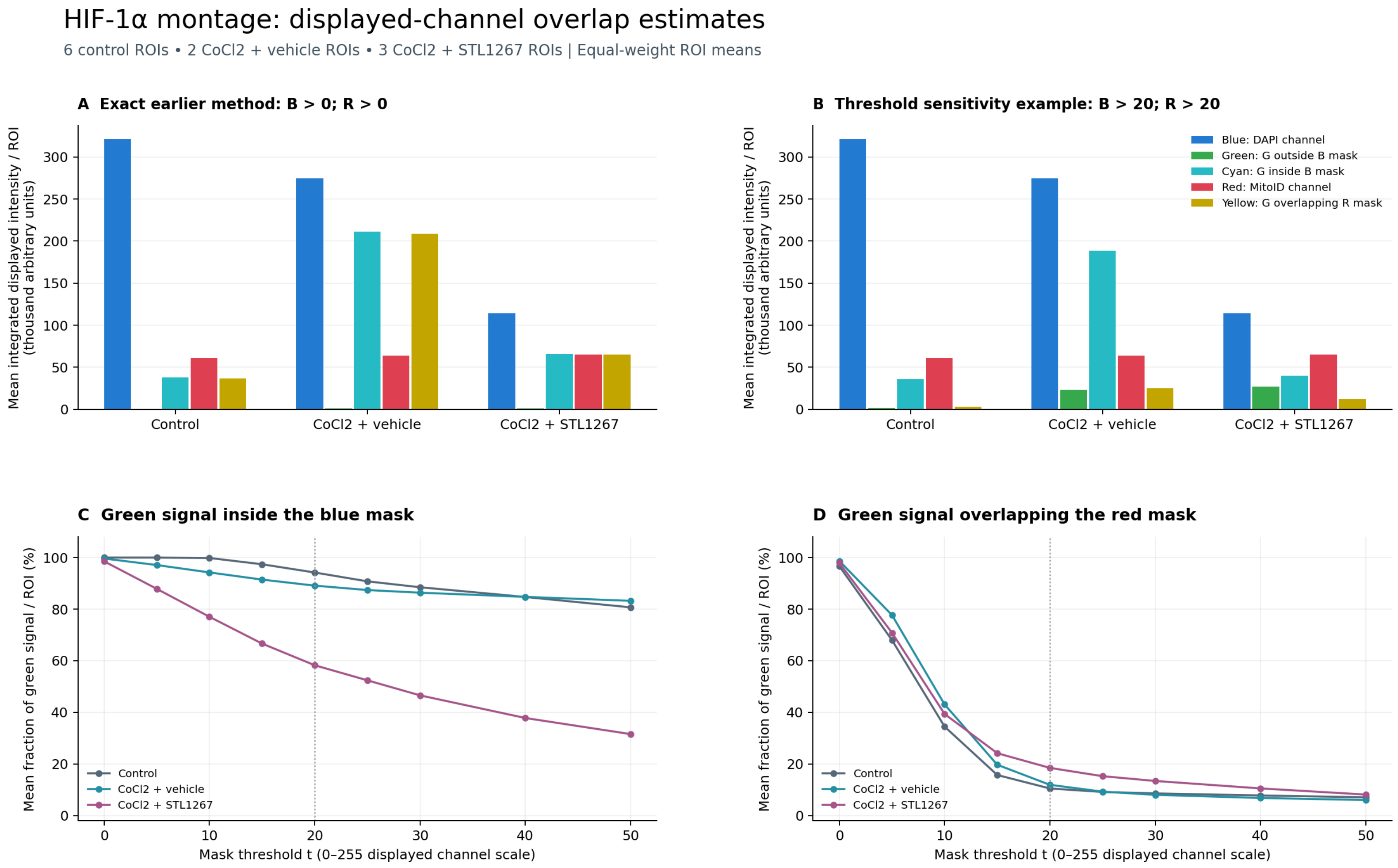


Blue and red are channel sums; cyan and yellow measure green intensity within blue/red masks. Categories overlap and must not be added.
Threshold 20 is a sensitivity example, not validated nuclear or mitochondrial segmentation. These are merged-image estimates from one field per condition.

| condition | threshold | n_rois | mean_blue_channel_sum | mean_green_total_sum | mean_green_outside_blue_sum | mean_green_inside_blue_sum | mean_red_channel_sum | mean_green_overlapping_red_sum | mean_inside_blue_percent | mean_outside_blue_percent | mean_overlapping_red_percent | mean_blue_mask_percent_of_roi | mean_red_mask_percent_of_roi | pooled_blue_channel_sum | pooled_red_channel_sum | pooled_green_total_sum | pooled_green_inside_blue_sum | pooled_green_outside_blue_sum | pooled_green_overlapping_red_sum |
|---|---|---|---|---|---|---|---|---|---|---|---|---|---|---|---|---|---|---|---|
| Control | 0 | 6 | 321282.6666666667 | 37869.833333333336 | 0 | 37869.833333333336 | 60892.5 | 36825.5 | 100 | 0 | 96.6341081 | 100 | 78.98346857790885 | 1927696 | 365355 | 227219 | 227219 | 0 | 220953 |
| CoCl2 + vehicle | 0 | 2 | 274411.5 | 211702.5 | 763 | 210939.5 | 63685 | 208963 | 99.63229196747659 | 0.36770803252340317 | 98.68639330618495 | 50.27758756443119 | 52.939749716398396 | 548823 | 127370 | 423405 | 421879 | 1526 | 417926 |
| CoCl2 + STL1267 | 0 | 3 | 114219.33333333333 | 66471.33333333333 | 926.3333333333334 | 65545 | 65102.666666666664 | 65182 | 98.53823083610189 | 1.4617691638981019 | 97.87694340575705 | 50.99057816098568 | 49.369585381051365 | 342658 | 195308 | 199414 | 196635 | 2779 | 195546 |
| Control | 5 | 6 | 321282.6666666667 | 37869.833333333336 | 4.666666666666667 | 37865.166666666664 | 60892.5 | 25755.166666666668 | 99.9904214 | 0.00957864 | 67.98791086082657 | 99.98712504784878 | 19.25596815286862 | 1927696 | 365355 | 227219 | 227191 | 28 | 154531 |
| CoCl2 + vehicle | 5 | 2 | 274411.5 | 211702.5 | 6168 | 205534.5 | 63685 | 164196 | 97.10237083137096 | 2.8976291686290416 | 77.69376944522094 | 35.15058861635361 | 32.620418460421135 | 548823 | 127370 | 423405 | 411069 | 12336 | 328392 |
| CoCl2 + STL1267 | 5 | 3 | 114219.33333333333 | 66471.33333333333 | 7712.666666666667 | 58758.666666666664 | 65102.666666666664 | 47392 | 87.86977631691184 | 12.130223683088163 | 70.8374381 | 25.251518053115017 | 22.66990858976837 | 342658 | 195308 | 199414 | 176276 | 23138 | 142176 |
| Control | 10 | 6 | 321282.6666666667 | 37869.833333333336 | 64.5 | 37805.333333333336 | 60892.5 | 12147.166666666666 | 99.84628214307698 | 0.1537178569230174 | 34.43687971075747 | 99.06774258904018 | 9.01122357 | 1927696 | 365355 | 227219 | 226832 | 387 | 72883 |
| CoCl2 + vehicle | 10 | 2 | 274411.5 | 211702.5 | 12305 | 199397.5 | 63685 | 91435.5 | 94.24032639686627 | 5.759673603133731 | 43.03391184471787 | 29.11357618666933 | 19.446918405473717 | 548823 | 127370 | 423405 | 398795 | 24610 | 182871 |
| CoCl2 + STL1267 | 10 | 3 | 114219.33333333333 | 66471.33333333333 | 14652.333333333334 | 51819 | 65102.666666666664 | 26201.333333333332 | 77.0440252 | 22.95597480643639 | 39.33323221866063 | 17.470266961941338 | 11.22196181592939 | 342658 | 195308 | 199414 | 155457 | 43957 | 78604 |
| Control | 15 | 6 | 321282.6666666667 | 37869.833333333336 | 997.6666666666666 | 36872.166666666664 | 60892.5 | 4950.166666666667 | 97.40562434034598 | 2.5943756596540246 | 15.691705152486458 | 65.20690193431768 | 6.07415678 | 1927696 | 365355 | 227219 | 221233 | 5986 | 29701 |
| CoCl2 + vehicle | 15 | 2 | 274411.5 | 211702.5 | 18275 | 193427.5 | 63685 | 41965.5 | 91.45989489887705 | 8.5401051 | 19.667245575253204 | 25.90381712292303 | 12.12600235029092 | 548823 | 127370 | 423405 | 386855 | 36550 | 83931 |
| CoCl2 + STL1267 | 15 | 3 | 114219.33333333333 | 66471.33333333333 | 21227.333333333332 | 45244 | 65102.666666666664 | 15861 | 66.65148353546157 | 33.34851646453843 | 24.14826870151977 | 13.983610141042343 | 7.42395177 | 342658 | 195308 | 199414 | 135732 | 63682 | 47583 |
| Control | 20 | 6 | 321282.6666666667 | 37869.833333333336 | 2008.5 | 35861.333333333336 | 60892.5 | 3128.3333333333335 | 94.20930305241734 | 5.790696947582652 | 10.472634615586756 | 47.901906981054445 | 5.153028593244919 | 1927696 | 365355 | 227219 | 215168 | 12051 | 18770 |
| CoCl2 + vehicle | 20 | 2 | 274411.5 | 211702.5 | 23319 | 188383.5 | 63685 | 25298.5 | 89.11944206135223 | 10.88055793864778 | 11.884775707329737 | 24.365279368011652 | 9.237219192919252 | 548823 | 127370 | 423405 | 376767 | 46638 | 50597 |
| CoCl2 + STL1267 | 20 | 3 | 114219.33333333333 | 66471.33333333333 | 26610.666666666668 | 39860.666666666664 | 65102.666666666664 | 12151.666666666666 | 58.25558813489288 | 41.74441186510712 | 18.45044614286881 | 12.204067551212011 | 6.097905635054474 | 342658 | 195308 | 199414 | 119582 | 79832 | 36455 |
| Control | 25 | 6 | 321282.6666666667 | 37869.833333333336 | 3025.8333333333335 | 34844 | 60892.5 | 2674.1666666666665 | 90.79036256581436 | 9.20963743 | 9.120687217820036 | 39.206612807807765 | 4.661634271789029 | 1927696 | 365355 | 227219 | 209064 | 18155 | 16045 |
| CoCl2 + vehicle | 25 | 2 | 274411.5 | 211702.5 | 26942.5 | 184760 | 63685 | 19615 | 87.41330644200389 | 12.586693557996117 | 9.246004077935115 | 23.42037113271398 | 7.844017986540523 | 548823 | 127370 | 423405 | 369520 | 53885 | 39230 |
| CoCl2 + STL1267 | 25 | 3 | 114219.33333333333 | 66471.33333333333 | 30461.333333333332 | 36010 | 65102.666666666664 | 10058.333333333334 | 52.41315318935228 | 47.58684681064772 | 15.24237684741739 | 11.051032748242932 | 5.203416412530781 | 342658 | 195308 | 199414 | 108030 | 91384 | 30175 |
| Control | 30 | 6 | 321282.6666666667 | 37869.833333333336 | 3731.3333333333335 | 34138.5 | 60892.5 | 2499.6666666666665 | 88.49860521278087 | 11.501394787219121 | 8.550511075269918 | 34.82070119320165 | 4.320444426775288 | 1927696 | 365355 | 227219 | 204831 | 22388 | 14998 |
| CoCl2 + vehicle | 30 | 2 | 274411.5 | 211702.5 | 29167 | 182535.5 | 63685 | 16977 | 86.38243750443137 | 13.617562495568635 | 8.005908258962029 | 22.849942751580585 | 6.927677378940277 | 548823 | 127370 | 423405 | 365071 | 58334 | 33954 |
| CoCl2 + STL1267 | 30 | 3 | 114219.33333333333 | 66471.33333333333 | 34311.666666666664 | 32159.666666666668 | 65102.666666666664 | 8855.666666666666 | 46.58427520342443 | 53.41572479657557 | 13.387384666951187 | 10.02531645880884 | 4.522806820825225 | 342658 | 195308 | 199414 | 96479 | 102935 | 26567 |
| Control | 40 | 6 | 321282.6666666667 | 37869.833333333336 | 4712.666666666667 | 33157.166666666664 | 60892.5 | 2261.6666666666665 | 84.72494050656108 | 15.275059493438933 | 7.7689247869647735 | 29.89784046049445 | 3.741715425234124 | 1927696 | 365355 | 227219 | 198943 | 28276 | 13570 |
| CoCl2 + vehicle | 40 | 2 | 274411.5 | 211702.5 | 32630 | 179072.5 | 63685 | 14345.5 | 84.7710755 | 15.22892454840299 | 6.7921272407340805 | 22.201282827309946 | 5.7983168235413345 | 548823 | 127370 | 423405 | 358145 | 65260 | 28691 |
| CoCl2 + STL1267 | 40 | 3 | 114219.33333333333 | 66471.33333333333 | 40270.666666666664 | 26200.666666666668 | 65102.666666666664 | 6839.333333333333 | 37.804809414356235 | 62.195190585643765 | 10.496890719175434 | 8.438872880909981 | 3.584516227074287 | 342658 | 195308 | 199414 | 78602 | 120812 | 20518 |
| Control | 50 | 6 | 321282.6666666667 | 37869.833333333336 | 5607.166666666667 | 32262.666666666668 | 60892.5 | 2036.6666666666667 | 80.71098763851141 | 19.28901236148859 | 7.035006343753099 | 27.199695173553398 | 3.2919508568448936 | 1927696 | 365355 | 227219 | 193576 | 33643 | 12220 |
| CoCl2 + vehicle | 50 | 2 | 274411.5 | 211702.5 | 35967.5 | 175735 | 63685 | 12635.5 | 83.22780558603168 | 16.77219441396832 | 5.990002527927484 | 21.528080142254716 | 5.129902233832571 | 548823 | 127370 | 423405 | 351470 | 71935 | 25271 |
| CoCl2 + STL1267 | 50 | 3 | 114219.33333333333 | 66471.33333333333 | 44561.666666666664 | 21909.666666666668 | 65102.666666666664 | 5179.333333333333 | 31.565417831893186 | 68.43458216810681 | 8.113535315792346 | 7.1058902 | 2.8590429835135027 | 342658 | 195308 | 199414 | 65729 | 133685 | 15538 |

Figure 2

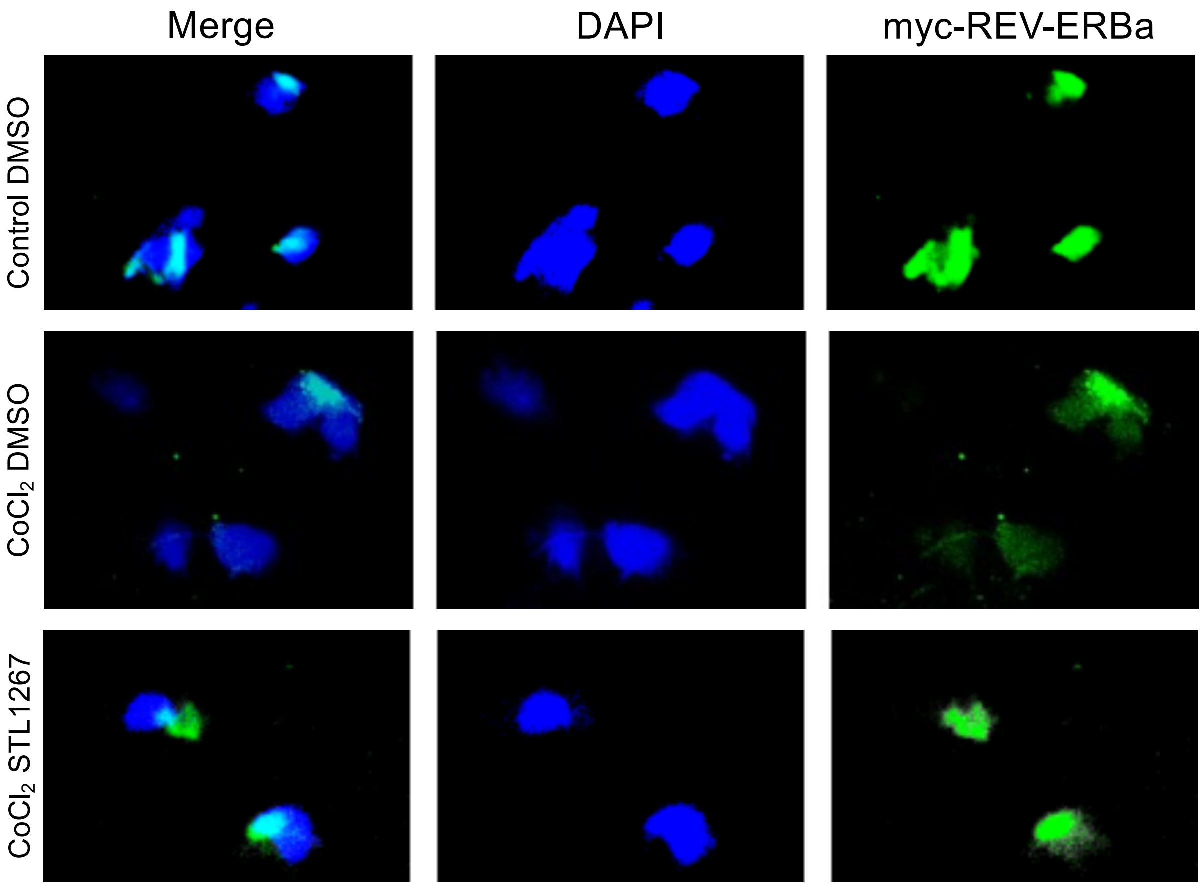

Merge
DAPI
myc-REV-ERBa
Control DMSO
CoCl2 DMSO
CoCl2 STL1267

### Control DMSO

#### Green signal inside and outside the blue region

Same pixel-overlap method as the preceding image • bars show the mean; symbols show each ROI

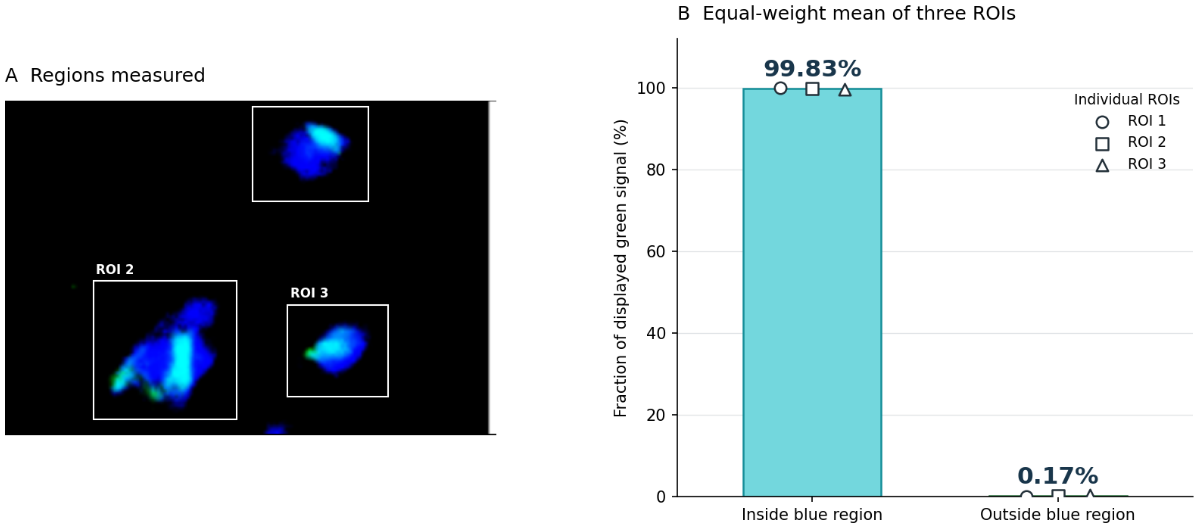


Three complete blue-stained regions were included. The partial blue object at the bottom edge was excluded.

Inside = blue > 0; outside = blue = 0 within each ROI. Percentages use summed displayed green values.
Any faint blue signal counts as inside; results depend on this mask definition and do not establish nuclear localization.

Merged RGB image estimate, not original-channel fluorescence quantification. No error bars or inferential statistics.
ROIs are blue-stained objects; the irregular lower-left region may contain unresolved adjacent cells.

| ROI | Inside blue | Outside blue |
|---|---|---|
| 1 | 99.997% | 0.003% |
| 2 | 99.823% | 0.177% |
| 3 | 99.675% | 0.325% |

CoCl2 + DMSO

#### Green signal inside and outside the blue region

Same pixel-overlap method as the preceding image • bars show the mean; symbols show each ROI

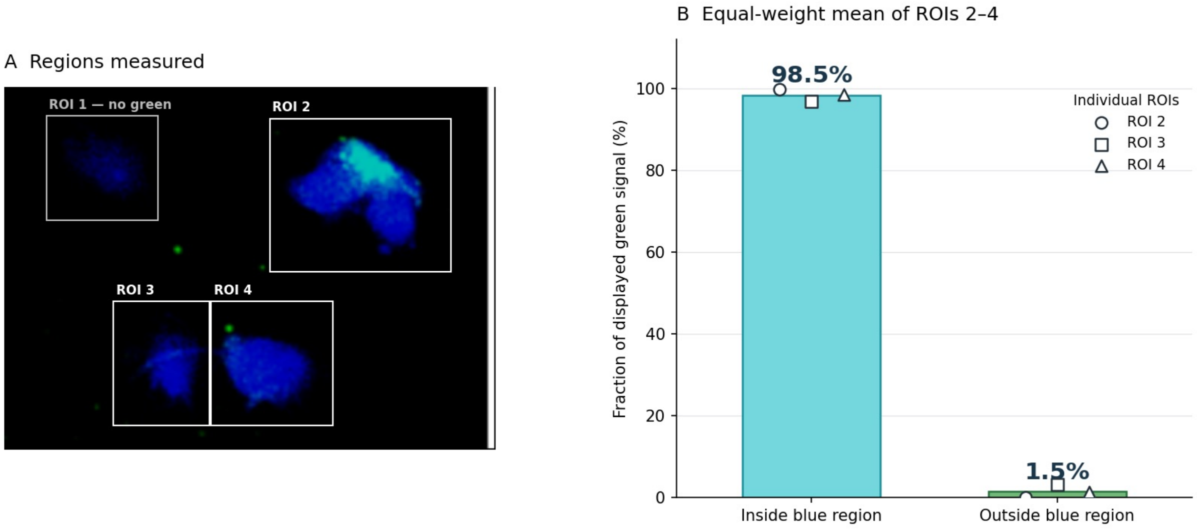


ROI 1 has zero green signal and is excluded from the percentage mean. ROI 2: 99.9% inside; ROI 3: 96.9%; ROI 4: 98.6%.

Inside = blue > 0; outside = blue = 0 within each ROI. Percentages use summed displayed green values.
Any faint blue signal counts as inside; results depend on this mask definition and do not establish nuclear localization.

Merged RGB image estimate, not original-channel fluorescence quantification. No error bars or inferential statistics.
ROIs are blue-stained objects; the irregular upper-right object may contain unresolved adjacent cells.

| Region | Inside blue | Outside blue |
|---|---|---|
| 1 | No green signal | Not calculated |
| 2 | 99.9% | 0.1% |
| 3 | 96.9% | 3.1% |
| 4 | 98.6% | 1.4% |

### CoCl2 + 1267

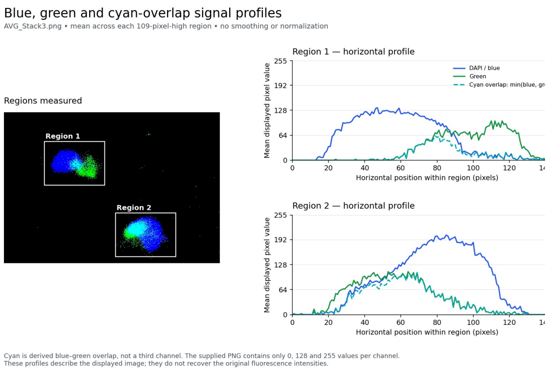


#### Green signal inside and outside the blue region

Two individual cells from one condition • bars show the mean; symbols show each cell

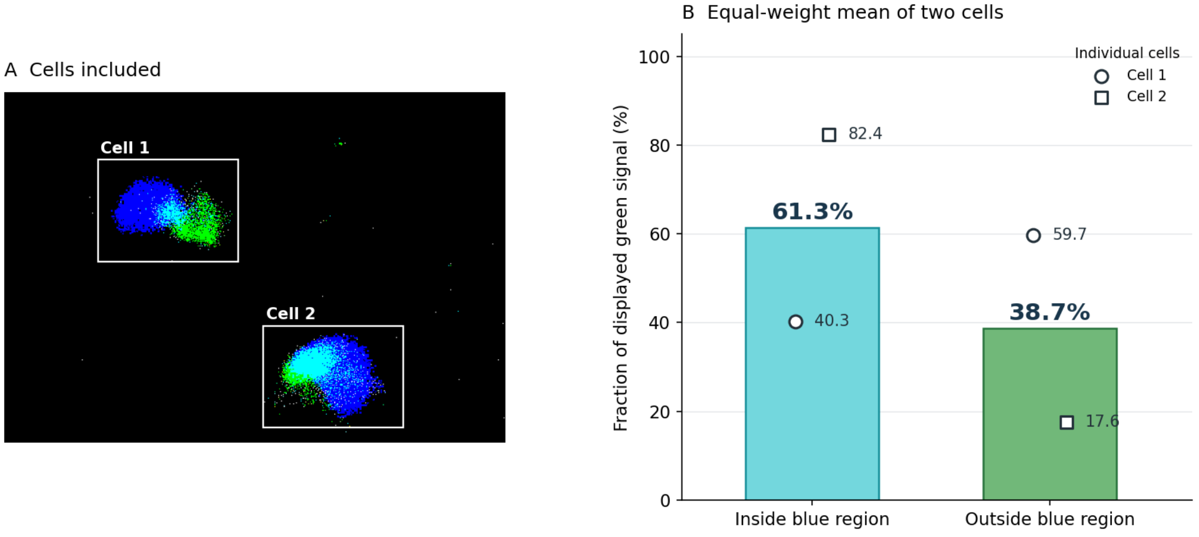


Inside = pixels with blue > 0; outside = pixels with blue = 0 within each outlined ROI.
Each cell's green signal was summed in both regions and expressed as a percentage of its total green signal.

Image-based overlap estimate from a saturated/thresholded RGB PNG. Not a validated nuclear–cytoplasmic measurement.
Two cells from one image are not independent biological replicates; no inferential statistics are shown.

| Cell | Inside blue | Outside blue |
|---|---|---|
| 1 | 40.3% | 59.7% |
| 2 | 82.4% | 17.6% |
